# The multifaceted roles of R2R3 transcription factor *Hl*MYB7 in the regulation of flavonoid and bitter acids pathways, development and biotic stress in hop (*Humulus lupulus* L.)

**DOI:** 10.1101/2022.10.03.510644

**Authors:** Ajay Kumar Mishra, Tomáš Kocábek, Vishnu Sukumari Nath, Ahamed Khan, Jaroslav Matoušek, Khaled M Hazzouri, Naganeeswaran Sudalaimuthuasari, Karel Krofta, Khaled M.A. Amiri

## Abstract

Hop (*Humulus lupulus*) biosynthesizes the highly economically valuable secondary metabolites, which include flavonoids, bitter acids, polyphenols and essential oils. These compounds have important pharmacological properties and are widely implicated in the brewing industry owing to bittering flavor, floral aroma and preservative activity. Our previous studies documented that ternary MYB-bHLH-WD40 (MBW) and binary WRKY1-WD40 (WW) protein complexes transcriptionally regulate the accumulation of bitter acid (BA) and prenylflavonoids (PF). In the present study, we investigated the regulatory functions of the R2R3-MYB repressor HlMYB7 transcription factor, which contains a conserved N-terminal domain along with the repressive motif EAR, in regulating the PF- and BA-biosynthetic pathway and their accumulation in hop. Constitutive expression of *HlMYB7* resulted in transcriptional repression of structural genes involved in the terminal steps of biosynthesis of PF and BA, as well as stunted growth, delayed flowering, and reduced tolerance to viroid infection in hop. Furthermore, yeast two-hybrid and transient reporter assays revealed that HlMYB7 targets both PF and BA pathway genes and suppresses MBW and WW protein complexes. Heterologous expression of *HlMYB7* leads to down-regulation of structural genes of flavonoid pathway in *Arabidopsis thaliana*, including a decrease in anthocyanin content in *Nicotiana tabacum*. The combined results from functional and transcriptomic analyses highlight the important role of *HlMYB7* in fine-tuning and balancing the accumulation of secondary metabolites at the transcriptional level, thus offer a plausible target for metabolic engineering in hop.

## Introduction

As sessile organisms, plants have evolved a diverse array of efficient and sophisticated mechanisms during their evolution to adapt flexibly under the influence of various hostile biotic and abiotic stresses (Yang et al., 2018). The response to such stimuli is orchestrated by cellular signaling components that lead to the activation of cellular responses, including the accumulation of a repertoire of secondary metabolites that represent the chemical interference between plants and their environment (Berini et al., 2018). Plant secondary metabolites are usually classified into several classes according to their chemical structure or biosynthetic pathway. Among them, phenylpropanoids represent a large class of secondary metabolites consisting of a diverse group of compounds with different biological functions (Bourgaud et al., 2001).

Phenylpropanoids play important roles in structural integrity, free radical scavenging, protection from UV light, modulation of developmental signaling cascades, pigmentation of co-evolutionary traits, root nodule formation, and internal regulation of plant cell physiology (Falcone Ferreyra et al., 2012). The shikimate pathway serves as a biosynthetic pathway for the production of the aromatic amino acids L-phenylalanine and as an entry point for the biosynthesis of phenylpropanoids with the participation of structural enzymes such as phenylalanine ammonia lyase (*PAL*), cinnamic acid 4-hydroxylase (*C4H*), 4-coumaryl:CoA ligase (*4CL*), chalcone synthase (*CHS*), chalcone isomerase (*CHI*), flavonol synthase (*FLS*), flavanone 3-hydroxylase (*F3H*), flavonoid 3’5’-hydroxylase (*F3’5’H*), dihydroflavonol-4-reductase (*DFR*), and anthocyanidin synthase (*ANS*) (Mouradov et al., 2014). The phenylpropanoid biosynthetic pathway is finely tuned and tightly regulated by the coordinated transcriptional control of genes encoding structural enzymes and the independent action of DNA-binding R2R3-MYB transcription factors (TFs) or the combinatorial interaction of R2R3-MYB, bHLH, and WD40 TFs (MBW complex) (Petroni and Tonelli, 2011; Biała and Jasiński, 2018). Although the coordinated interaction of the MBW complex activates transcription, but the specificity of the complex and the determination of specific pathway regulation is regulated by the involvement of the corresponding R2R3-MYB TFs (Cavallini et al., 2015). Furthermore, a single R2R3-MYB gene was discovered to function as both a repressor and activator of genes, and the redundancy, dual function, and opposing regulatory roles in common target genes by R2R3-MYBs enable fine-tuning of spatiotemporal transcriptional regulation of plant secondary metabolic pathways (Bhargava et al., 2010; Reddy et al. 2017). In plants several R2R3-MYBs have been characterized as activators or repressors of genes involved in the flavonoid, flavonol, phenylpropanoid and anthocyanin pathways (Reddy et al., 2017). For instance, three closely related MYB-TFs, namely AtMYB12, AtMYB11, and AtMYB111, act as activators of the genes *CHS, CHI, F3H*, and *FLS*, which in turn regulate flavonols content in *Arabidopsis thaliana* (Mehrtens et al., 2005; Tracke et al., 2007). Similarly, TFs GmMYB12B2 and GmMYB176 positively regulate the expression of CHS8 and CHS, which are involved in isoflavonoid biosynthesis in soybean (Yi et al., 2010). In addition to their role as activators, the two distinct groups of MYB TFs, namely R3-MYBs and R2R3-MYB, which contain one or two repeats of the MYB domain region, function as negative regulators in phenylpropanoid and flavonoid biosynthesis (Dubos et al., 2008; Matsui et al., 2008). The R2R3-MYB TFs have been divided into 22 subgroups based on the presence of specific conserved regions in the C-terminus, with the majority of MYB repressor proteins belonging to subgroup 4, which can be further classified into general phenylpropanoid, lignin, and flavonoid groups based on the target pathway (Liu et al., 2015). These MYB repressors are characterised by the presence of repression domains in the C-terminal region, such as the C1 domain (GIDP motif), the C2 domain (EAR repression motif) defined by the consensus sequence patterns of either LxLxL or DLN xxP (Jin et al., 2000; Kagale and Rozwadowski, 2011), and the C4 domain with the conserved dFLGL and LDF/YRxLEMK amino acid signatures (Kagale and Rozwadowski, 2011). A growing body of research suggests that competition between MYB activators and MYB repressors for binding cofactors or DNA cis-elements is a mechanism underlying repressive activity. The flavonoid MYB repressors can either competitively influence the formation of the MBW activation complex by binding bHLH coactivators or convert them into repressive complexes via their EAR motif (Albert et al., 2014; Yoshida et al., 2015). Strawberry FaMYB1 (Aharoni et al. 2001), petunia PhMYB27 (Albert et al., 2014) and ZmMYB31, ZmMYB42 from maize (Fornalé et al., 2010) represent this class of TFs.

Hop (*Humulus lupulus* L.) is a diploid, dioecious, perennial climbing plant in the Cannabaceae family, grown exclusively for the brewing industry, but also important as a feed supplement in livestock and for medicinal purposes (Mishra et al. 218). The female inflorescences of hop plants (cones) contain metabolically active glandular trichomes known as “lupulin glands” that serve as molecular secretory factories for the synthesis and accumulation of specialized secondary metabolites such as prenylated flavonoids (xanthohumol and desmethylxanthohumol), bitter acids (humulone or α-acid and lupulone or β-acid), essential oils, and terpenoids (Roberts et al., 2004). Several lupulin gland-specific structural genes encoding key enzymes of the prenyflavonoid and bitter acid pathways, such as chalcone synthase (*HlCHS*_H1) (Matoušek et al., 2006), prenyltransferase1 (*HlPT-1*) (Tsurumaru et al., 2010) and O-methyltransferase 1 (*OMT1*) and their transcriptional regulators, which belong to the MYB, bHLH, WDR, and WRKY families, are well documented in hop (Matoušek et al., 2012; Kocábek et al., 2018; Mishra et al., 2018). Parallel activation of the *CHS*_H1 promoter was shown to be driven by highly organized ternary MBW complexes (HlMYB2/HlbHLH2/HlWDR1), whereasa binary complex of *Hl*WRKY1 and *Hl*WDR1 (WW) was reported to activate the structural genes of the terminal stages and the ternary MBW complex of the prenylflavonoid (PF) and bitter acids (BA) biosynthetic pathways (Matoušek et al. 2012; Matoušek et al., 2016; Mishra et al., 2018). However, the presumed role of MYB repressors in terms of their ability to affect specific structural genes and MBW/WW protein complexes of the biosynthetic pathways PF and BA is still enigmatic. In the present study, we successfully isolated and cloned the cDNA of the *HlMYB7* gene based on primary sequence homology with the R2R3-MYB repressor gene AtMYB4 of Arabidopsis. Our detailed investigation using multidisciplinary approaches sheds new light on the coordinated involvement of HlMYB7 TF in fine-tuning the regulation PF and BA of biosynthesis, growth, and biotic stress responses in hop.

## Results

### Isolation and sequence analysis of the R2R3-MYB C2 repressor *HlMYB7* in hop

A cDNA clone with an open reading frame sequence of 798 bp encoding a putative R2R3-MYB protein with 265 amino acid residues and a predicted molecular weight and calculated isoelectric point of 29.56 kDa and 7.43 kDa, respectively, was isolated from hop by RT-PCR. The isolated sequence was designated *HlMYB7* and submitted to GenBank under accession number FR873650.1. The deduced amino acid sequence consists of the highly conserved R2 and R3 MYB DNA-binding sites at the amino-terminal end, while the C-terminal domain is highly diverse and contains three conserved protein motifs, the C1 motif (LlsrGIDPxT/SHRxI/L), the C2 motif (pdLNLD/ELxiG/S), and the C4 motif (FLGLx4-7V/LLD/GF/ YR /Sx1LEMK) (Figure 1a), which are mainly found in AtMYB4-like transcription inhibitors (Kagale and Rozwadowski, 2011). Phylogenetic analysis of 19 plant MYB proteins associated with different functions revealed that HlMYB7 belongs to the same cluster as VvMYB4a, MnMYB308, and CsMYB308-like (Figure 1b), which are known to play important roles as transcriptional repressors in phenylpropanoids biosynthesis (Cavallini et al., 2015). The HlMYB7 protein sequence has an overall sequence similarity of 90% and 78% with its closest neighbors, the CsMYB308 and MnMYB308 proteins, respectively (Figure 1b). Overall, phylogenetic and sequence analysis suggests that *HlMYB7* is a novel gene belonging to the R2R3-MYB TF family and distinguished by the presence of a C2 repression motif that may be involved in the regulation of PF and/or BA synthesis in hop.

**Figure 1.**
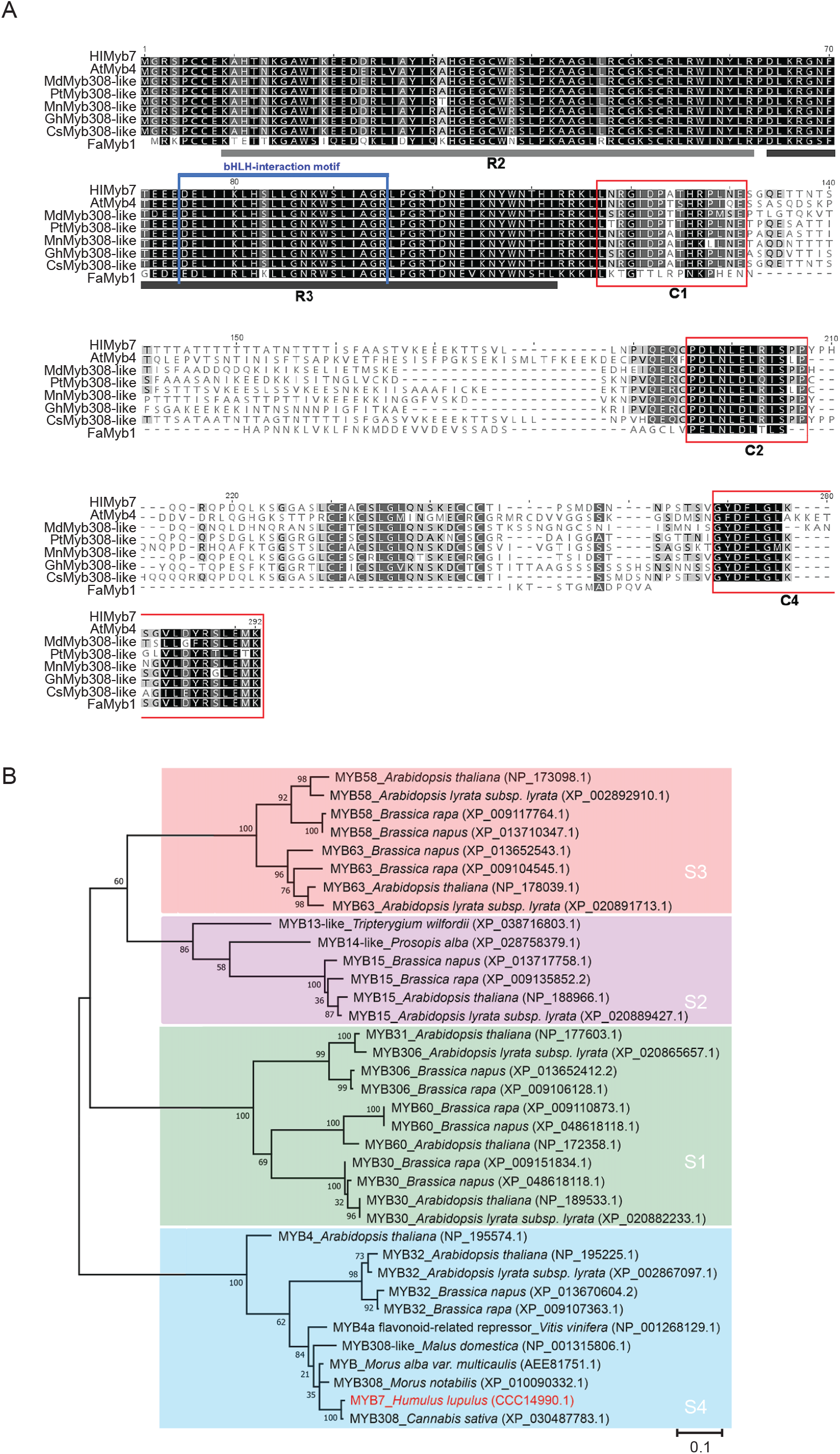
Sequence alignment and phylogenetic analysis of HlMYB7 protein. (**A**) Alignment of the deduced amino acid sequences of HlMYB7 and selected R2R3-MYB regulators of phenylpropanoid biosynthesis from other plant species AtMYB4 (*Arabidopsis thaliana*), MdMYB308 (*Malus domestica*), PtMYB308 (*Phaeodactylum tricornutum*), MnMYB308 (*Morus notabilis*), GhMYB308 (*Gossypium hirsutum*), CsMYB308 (*Camellia sinensis*), FaMYB1 (*Fragaria ananassa*). The R2 and R3 MYB domains are underlined and refer to two repeats of the MYB DNA-binding domain of MYB protein, whereas C1, C2, and C4 motifs outside of R2R3 MYB domain are indicated by red boxes. The numbers refer to the positions of the amino acid sequences. The identical amino acids are marked in black and the blue boxes indicate the bHLH interaction domain. (**B**) Phylogenetic relationships between HlMYB7 and R2R3-MYB transcription factors from other plant species. Amino acid sequences retrieved from the GenBank database were aligned using Clustal W, and the phylogenetic tree was constructed with MEGA 11 using the neighbor-joining method and the p-distance model. The scale bar represents 0.1 substitution per site, and the numbers next to the nodes are bootstrap values from 1,000 replicates. The accession numbers of the retrieved R2R3-MYB protein sequences from the GenBank database are embedded in the diagram.

### Tissue*-*specific and transcriptional repressor activity of *HlMYB7* on PF and BA biosynthetic genes

Spatiotemporal expression of *HlMyb7* TF was monitored in seven different hop tissues-roots, leaves, petioles, flowers, bracts, pollen, and lupulin gland stems-at different developmental stages by qRT-PCR (Figure 2a). The result showed that the transcription level of *HlMYB7* was significantly higher in lupulin glands than in other tissues where expression was comparable and lowest (∼7-10% of lupulin gland expression), suggesting its important role in flavonoid biosynthesis.

**Figure 2.**
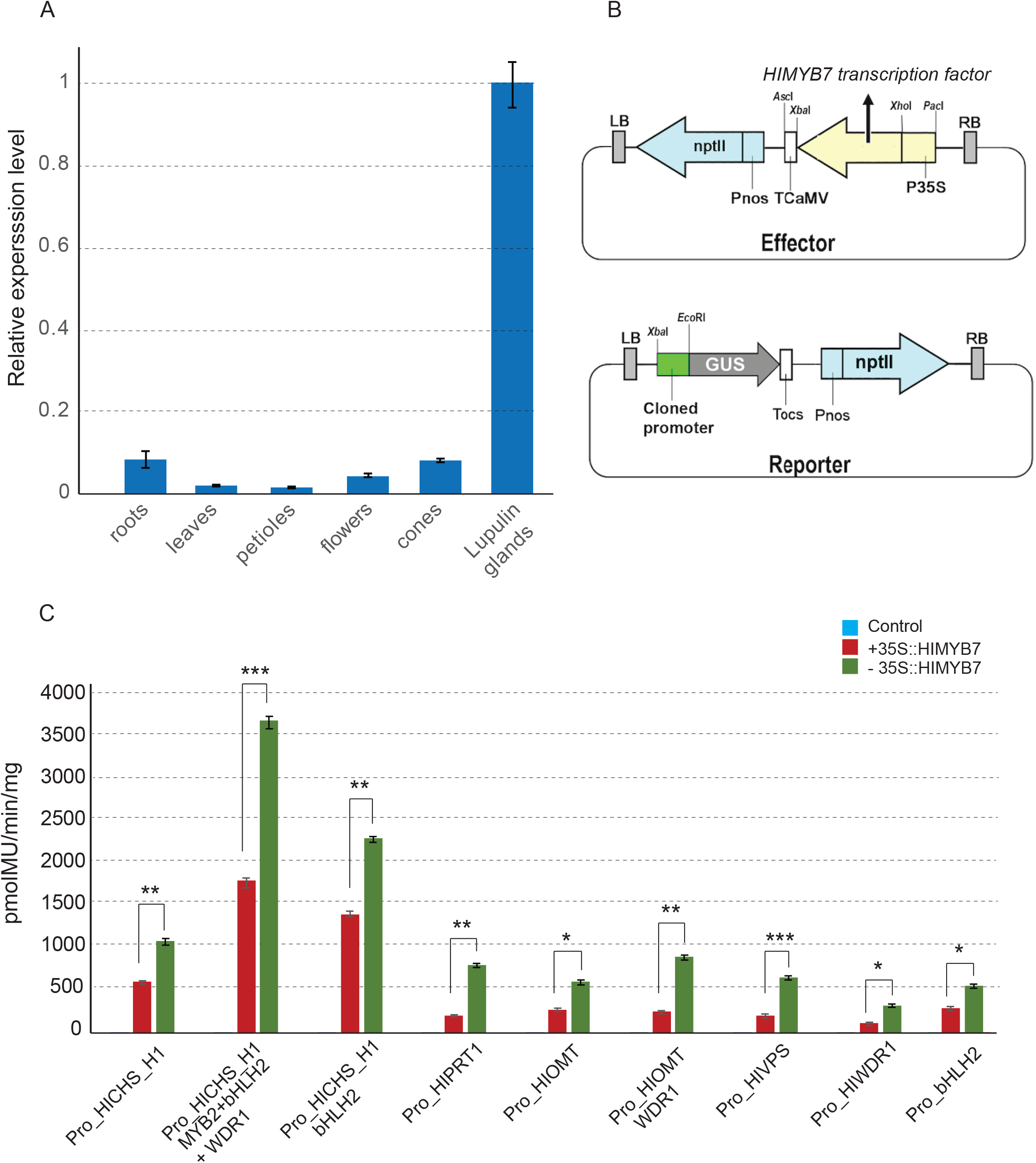
Expression profile and transcriptional activity of *HlMYB7*. (**A**) RT-qPCR analysis of the relative expression level of *HlMYB7* in different tissues of wild-type hop. Expression levels are normalized relative to those of *GAPDH*, which was used as an internal control. Error bars represent ± SD of three biological replicates. (**B**) Schematic diagrams illustrate the effector and reporter used in the transcriptional activity assay. The gene *GUS* was driven by the cloned promoter, whereas *HlMYB7* or the regulators were driven by the 35S promoter. (**C**) Transcriptional repressive effect of *HlMYB7* on the promoters of genes related to the prenylflavonoid and bitter acid pathways with and without cofactor in transiently transfected leaves of *Nicotiana benthamian*. Infiltrated leaves of *N. benthamiana* with a promoter-GUS reporter construct and a reporter construct lacking *HlMYB7* were used as controls. Each column represents the mean value ±SD of three independent experiments. Asterisk indicates statistically significant differences, *significant at p<0.05); **significant at p<0.01, *** significant at p<0.001.

To investigate the transcriptional activity of *Hl*MYB7 in regulating the genes encoding the structural enzymes of the PF and BA pathways, we used a transient expression assay system in leaves of *N. benthamiana* with promoters of the genes encoding the enzymes of the final stages of the biosynthesis of PF (*CHS*_H1, *OMT, PT1*) and BA [valerophenone synthase (*VPS*)] with (either individually or in combination) or without transcriptional cofactors (*Hl*MYB2, *Hl*bHLH2, *Hl*WDR1, and *Hl*WRKY1) using reporter and effector constructs (Figure 2b). The results showed that the β-glucuronidase activity of the promoters of the structural genes *CHS_H1, OMT* and *PT1* was reduced by approximately 1.5-to 4-fold when *HlMYB7* was used alone, but the repression of these promoters was significantly increased when 35S:*HlMYB2*, 35S:*HlbHLH2*, 35S:*HlWDR1* were coexpressed simultaneously compared with the coexpression of 35S:*HlbHLH2* (Figure 2c). It is noted that the activity of the promoter of transcriptional cofactors (*Hl*bHLH2 and *Hl*WDR1) was moderately suppressed upon infiltration with 35S:*HlMYB7* (Figure 2c). Additionally, the suppression of *HlOMT* was considerably enhanced when *HlWDR1* regulators were co-transformed. Taken together, these results suggest that HlMYB7 can repress the expression of structural genes associated with the PF and BA pathways and requires HlbHLH2 and HlWDR1 as regulatory proteins to enhance the ability of transcriptional repression at the promoters of the *CHS*_H1 and *OMT* genes.

### HlMYB7 is a nucleus-localized protein and suppresses the activation effect of MBW and WW on the promoters of *HlCHS*_H1 and *HlOMT1*, respectively

To investigate subcellular localization, GFP was fused to the C-terminus of *HlMYB7* and introduced into the epidermal cells of *Nicotiana benthamiana*. GFP fluorescence of the HlMYB7-GFP fusion protein was detected exclusively in the nuclei, whereas fluorescent signals of the control were observed in whole cells (Figure 3a), indicating that HlMYB7 is localized in the nucleus. Considering the presence of the conserved bHLH interaction motif in the HlMYB7 protein, the Y2H assay was primarily performed to detect the interaction between HlMYB7 and HlbHLH2 and subsequently with other proteins of the MBW and WW complexes involved in biosynthesis PF and BA.. Yeast cells cotransformed with combinations of HlMYB7-BD and HlbHLH2-AD or HlMYB7-BD and HlWDR1-AD were able to grow on selective media lacking His and Ade, indicating a strong interaction of HlMYB7 with HlbHLH2 and HlWDR1 and presumably contributing to its activity as repressors. This conclusion was also confirmed by the formation of blue colonies in the corresponding yeast cells as determined by the β-galactosidase assay (Figure 3b)

**Figure 3.**
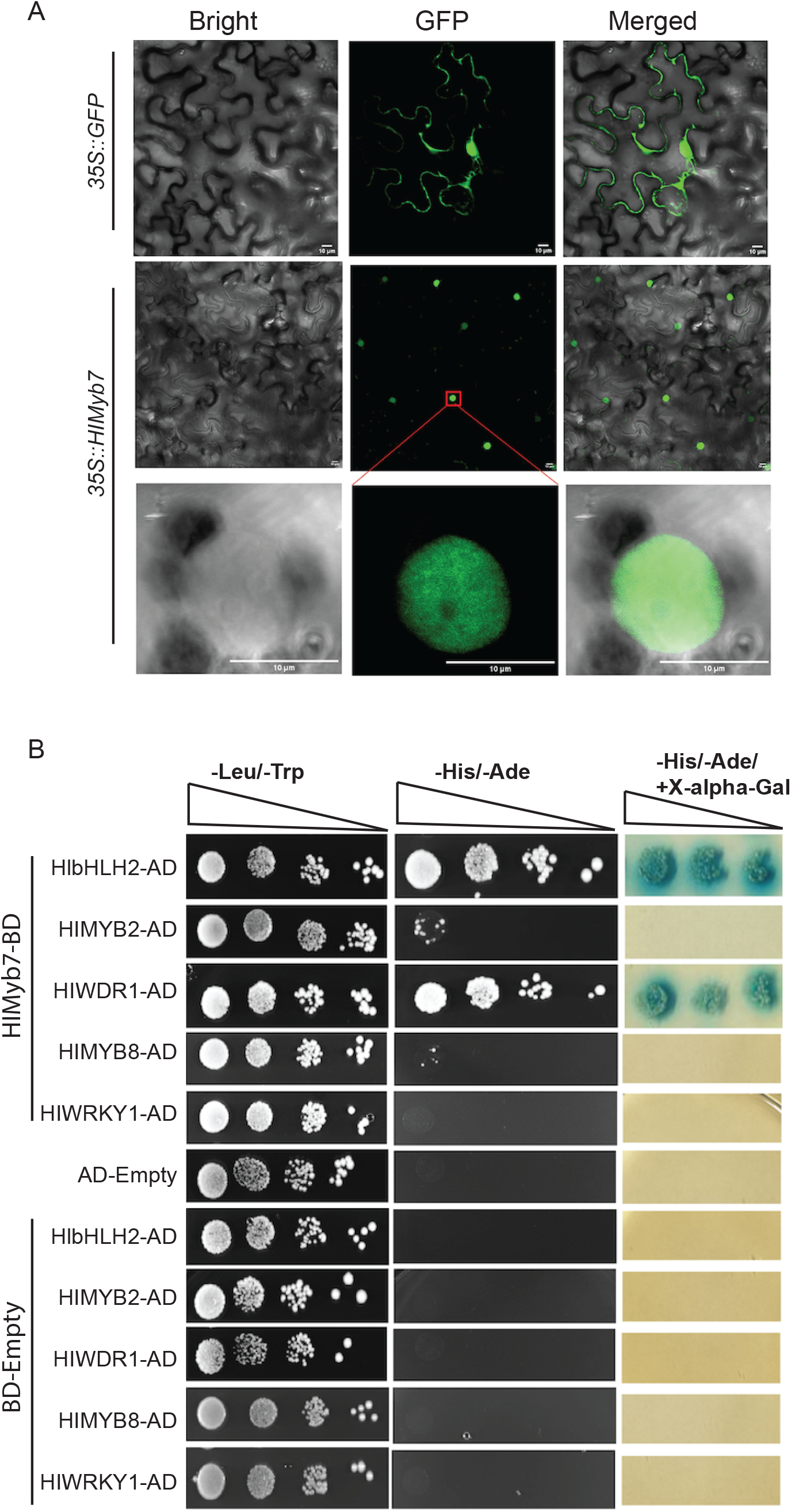
Subcellular localization and Yeast two-hybrid assay of *HlMYB7* transcription factor. (**A**) Confocal images showing nuclear localization of HlMYB7-GFP fusion protein (middle), a 100× magnified view of a single nucleus expression HlMYB7-GFP (bottom) and nuclear and cytomembrane localization of GFP alone (top) in leaf epidermal cells of *N. benthamiana*. Scale bars = 10 μm. (**B**) Yeast two-hybrid (Y2H) analysis of HlMyb7 interaction network with the transcription factors HlbHLH2, HlMYB2, HlWDR1, HlMYB8, and HlWRKY1. Co-transformation of bait and prey vectors for each combination was cultured on both nonselective media (SD/-Trp/-Leu/+Ade/+His) and selective media (SD/-Trp/-Leu/-Ade/-His) and with X-α-Gal (SD/-Trp/-Leu/-His/X-α-Gal). For each interaction, the control was combined with empty prey and bait and empty bait with the prey construct.

### Overexpression of *HlMYB7* leads to changes in morphological features and decreased biosynthesis and accumulation of flavonoids and bitter acids in hop

To gain insight into the range of biological functions regulated by the *HlMYB7* gene, we performed agroinfiltration of nodal segments of *in vitro*-grown hop plants with a binary vector pLV07 containing 35S:*HlMYB*7 overexpression constructs (Figure S1a). Of the 215 regenerated antibiotic-resistant shoots, 18 plants successfully rooted on medium supplemented with kanamycin. Southern blot and PCR amplification with specific *nptII* primers confirmed seven successful independent transformation events (Figure S1b, c), whereas qRT-PCR confirmed differential expression of the transgene (*HlMYB7*) (10- to 50-fold) in Hl-HM7T transformants (Figure S1d). Based on these analyses, three independent Hl-HM7T lines with one lower and two higher transgene expressions (Hl-HM7T-1, Hl-HM7T-5, and Hl-HM7T-11) were selected and grown in the greenhouse alongside control hop plants. The Hl-HM7T lines exhibited growth stunting (Figure S1e), slightly later flowering (delayed four months), and without significant effects on cone dry weight, lupulin gland density, and chlorophyll content compared with control plants under normal greenhouse conditions (Table S2).

To investigate putative transcriptional relationships between *HlMYB7* and terpenophenolics (PF and BA) biosynthesis-related genes, the expression levels of key structural and regulatory genes were measured by qRT-PCR in lupulin glands isolated from cone tissue. The expression levels of the structural genes *PAL, CHS*_H1, *PT1, OMT, VPS, F3H, DFR*, and *FLS* were suppressed in Hl-HM7T lines compared with control hop plants, suggesting that HlMYB7 has repressive potential on the key enzymatic players of the flavonoids and BA biosynthetic pathways (Figure 4a). Strong down-regulation was observed for *CHS_H1* and downstream structural genes of the PF and BA pathways, further highlighting the repressive potential of HlMYB7 by disrupting the interaction of MBW-tripartite and WW-bipartite complexes with their structural genes.

**Figure 4.**
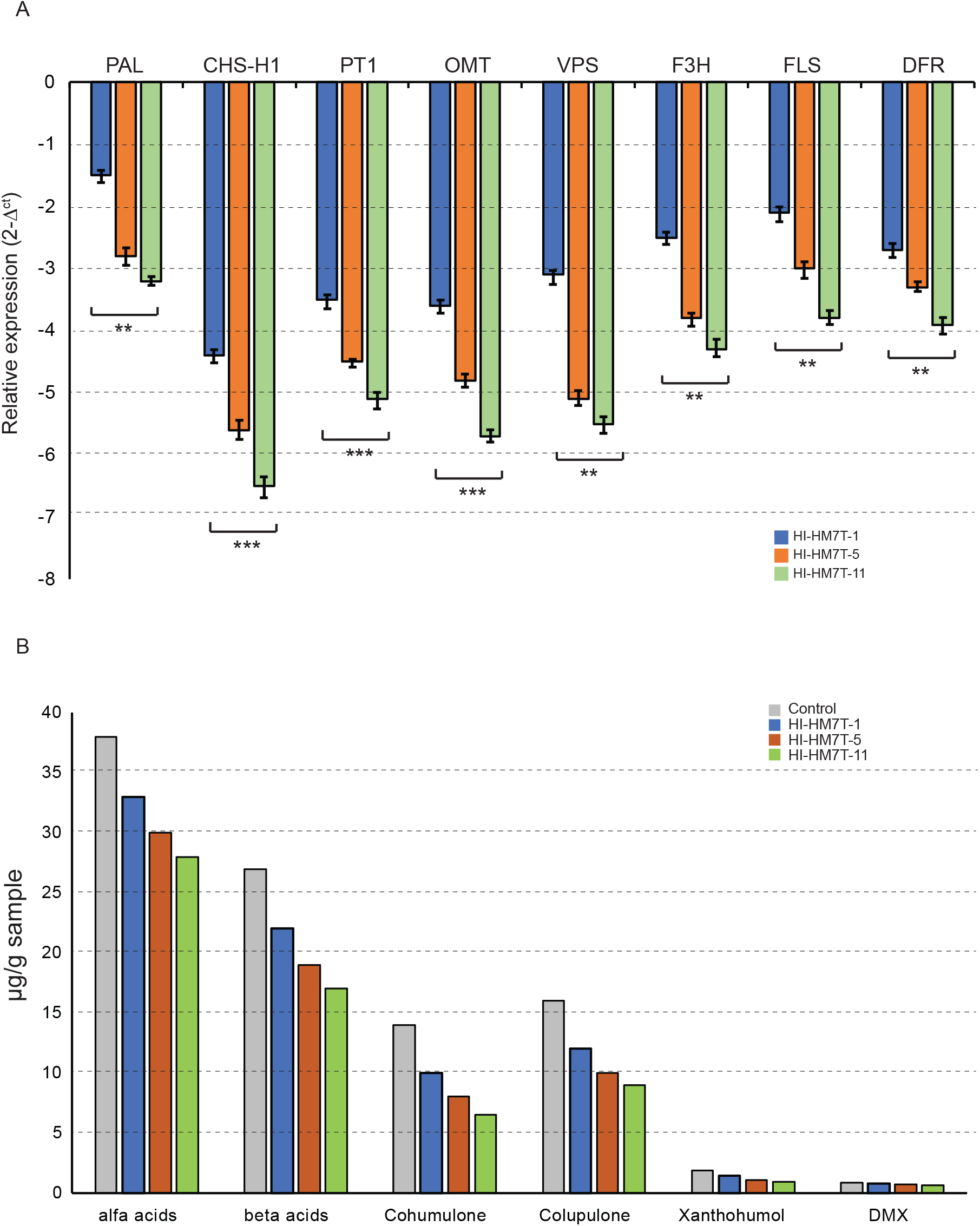
Functional analysis of HlMYB7 repressors in hop. (**A**) RT-qPCR analysis of the relative expression levels of endogenous prenylflavonoid and bitter acids biosynthetic genes in a in lupulin glands isolated from cone tissues of WT and MYB7 overexpression lines (Hl-HM7T-1, Hl-HM7T-5 and Hl-HM7T-11) of hop. PAL: phenylalanine ammonia-lyase; CHS_H1: chalcone synthase isoform 1; PT1: prenyltransferase 1; OMT: O-methyltransferases; VPS: valerophenone synthase; F3H: flavanone 3-hydroxylase; FLS: Flavonol synthase, and DFR: Dihydroflavonol 4-reductase. qRT-PCR analyses were normalised using GAPDH as an internal control gene. The fold change of each gene was calculated using the 2^−ΔΔCT^ method. Each column represents the mean value ±SD of three independent experiments. Asterisk indicates statistically significant differences, *significant at p<0.05; **significant at p<0.01, *** significant at p<0.001. (**B**) HPLC analysis of methanolic extracts of lupulin glands isolated from dried cones of WT and Hl-HM7T lines. Quantification (µg gm^-1^) of lupulone derivatives (α-acids and β-acids), prenylated acylphloroglucinols (cohumulone and colupulone) and prenylflavonoids [desmethylxanthohumol (DMX) and xanthohumol (X)] was performed using their respective working standards.

Nevertheless, comparative analysis of the HPLC-UV profiles showed a significant reduction in the content of xanthohumol, alpha and beta acids in the cones of the HM7T lines compared to the control hop plants (Figure 4b). These results correlate the overexpression of *HlMYB7* with the down-regulation of structural gene expression in HM7T lines, supporting the repressive role of HlMYB7 in controlling the enzymatic steps of the PF and BA pathways.

### Overexpression of *HlMYB7* effects the transcriptome landsacpe in HM7T lines

Overexpression of HlMYB7 results in significant morphological changes and delayed flowering, indicating widespread changes in gene expression at the molecular level. In this context, we performed comparative transcriptome profiling using RNAseq libraries generated from leaf tissues of three independent HM7T lines and control plants along with their technical replicates to assess the impact of *HlMYB7* overexpression on global changes in gene expression in hop. The high-throughput sequencing run generated over 36 and 60.35 million high-quality paired-end raw reads in control and HM7T lines, respectively. After removing adaptor sequences, low-quality reads, and trimming sequences that contained homopolymers longer than 8 bp, a total of 33 and 49 high-quality filtered reads were obtained for the control and HM7T lines, respectively.

Differential expression analysis revealed that 778 unigenes were modulated in HM7T lines compared with control, with 288 significantly upregulated and 490 significantly downregulated (Supplemental Table S3). Among the differentially expressed genes (DEGs), enzyme genes involved in the biosynthesis of flavanoids, flavonols, BA, anthocyanins, and protoanthocyanidins showed reduced expression in HM7T lines. Unexpectedly, *CONSTANS-LIKE* (*COL*) and CCT domain-containing genes that play important roles in photoperiod signalling and flowering time modulation (Chen et al., 2021) exhibited reduced expression. Furthermore, we also noticed that DEGs involved in host immune responses such as patatin-like protein 2 (La Camera et al., 2009), pathogenesis-related protein 4 (PR-4), PR-5 (Guevara-Morato et al., 2010), including TIFY proteins (Zhang et al., 2020), β-glucanases, chitinase-like (CTL) proteins (Perrot et al., 2022), leucine-rich repeat receptor kinases (Torii, 2004) which regulate a variety of developmental and defence-related processes, were downregulated (Supplemental Table S3). qRT-PCR analysis of some of these genes, including ten randomly selected DEGs (five up-regulated and five down-regulated genes) was consistent with the results of the RNA-seq data (Supplemental Figure S2), demonstrating the reproducibility of the transcriptome data.

Gene Ontology (GO) enrichment analysis of the 162 annotated up-regulated genes showed that cellular processes, response to stimuli, and response to chemicals were significantly enriched, whereas molecular functional categories related to plant development (such as cell wall modification, wax biosynthesis, and hormonal regulation), lipid metabolism, and catabolic processes were overrepresented among the 298 annotated down-regulated genes (Supplemental Figure S3a). Strikingly, KEGG-based enrichment analysis annotated downregulated unigenes in the categories of biosynthesis of secondary metabolites and biosynthesis of phenylpropanoids (Supplemental Figure S3b). Furthermore, an unbiased systematic overview of the involvement of DEGs in biological processes through MapMan analysis further cinfirmed the GO and KEGG enrichment analyses (Supplemental Figure S3c). MapMan analysis revealed suppression of genes related to the mevalonate metabolic pathway, hormone signalling, cell wall metabolism, and secondary metabolite biosynthesis pathway, which was consistent with the GO and KEGG enrichment analyses. These results were strongly correlated with stunted growth, delayed flowering, and a decrease in PF and BA content in HM7T lines.

### *HlMYB7* negatively regulates resistance against AFCVd-infection in hop

Compared with control hop, overexpression of *HlMYB7* suppressed a number of genes involved in immune responses in plants. To investigate the role of HlMYB7 in the defense responses against viroid diseases, the successfully acclimated three-month-old, well-rooted three transgenic Hl-HM7T lines (2, 13, and 17) and control plantlets were biolistically inoculated with an infectious dimeric construct of the cDNA of AFCVd (Figure 5a) and examined for disease severity and incidence of infection after dormancy (270 dpi) by dot-blot analysis, RT-PCR, and ssRT-qPCR. After the initial dormancy period, control plants of AFCVd-infected hops showed modest morphological changes with no apparent effects on plant growth and development, whereas the three AFCVd-infected Hl-HM7T lines exhibited severe developmental defects with leaf malformations (Figure 5b). The RT-PCR product of approximately 266 bp was observed in samples positive for AFCVd infection in dot-blot analysis (Figure 5c). Intriguingly, dot-blot hybridization and ssRT-qPCR analysis revealed a higher level of AFCVd (+) strands in Hl-HM7T lines compared with control hop plants artificially infected with AFCVd (Figure 5d, e). Expression of HlMYB7 was increased 15- to 25-fold in AFCVd-infected control hops, further confirming its negative regulatory role in viroid infection.

**Figure 5.**
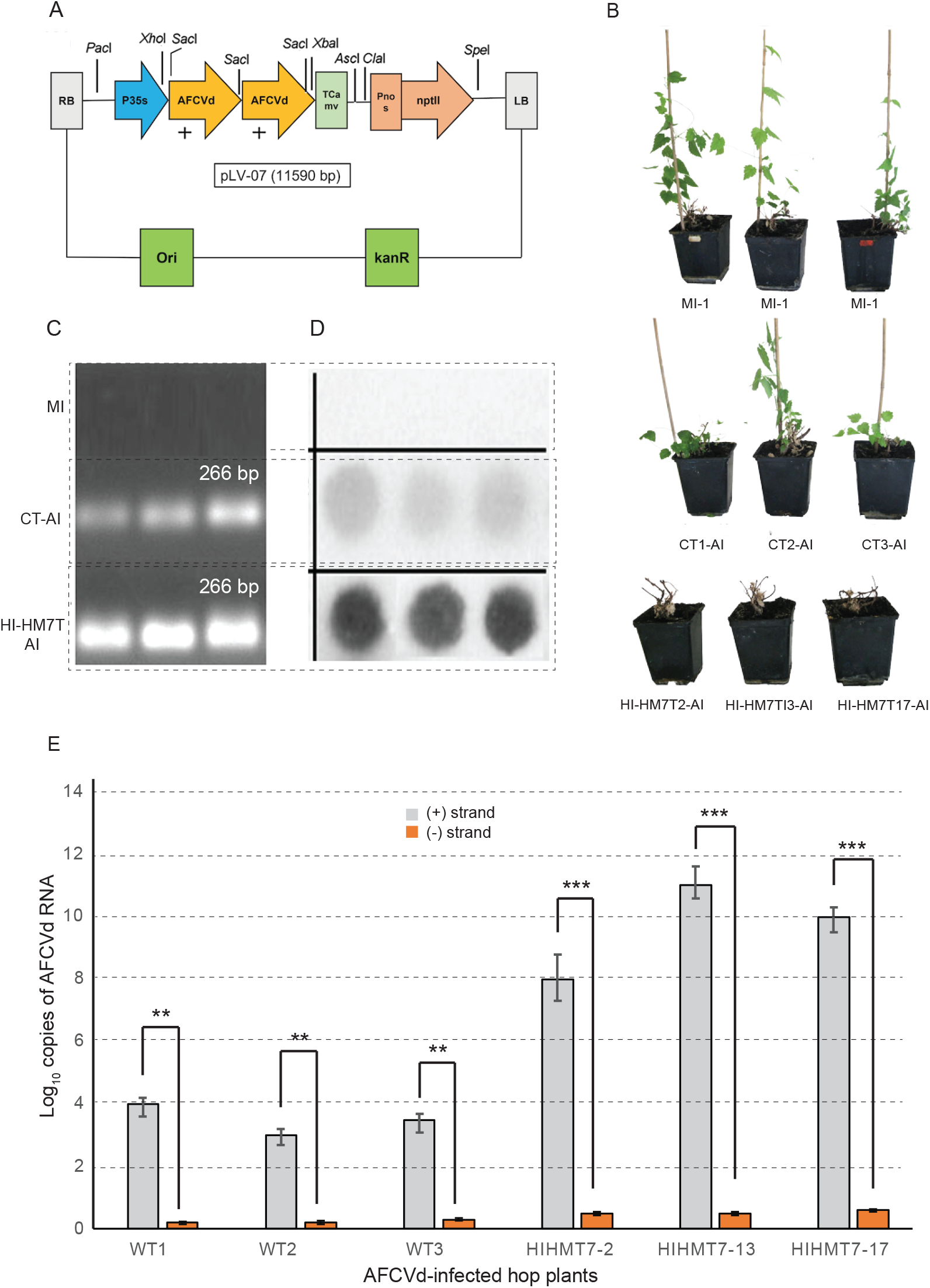
Construction of a dimeric cDNA construct for apple fruit crinkle virus (AFCVd) and detection and quantification of AFCVd in leaf tissue of WT and MYB7 overexpression lines (Hl-HM7T-1, Hl-HM7T-5, and Hl-HM7T-11) of hop. (**A**) Schematic representation of a plasmid containing the AFCVd (+) dimer generated by cDNA cloning at the *Sac*I restriction site. ori: origin of replication; kanR: kanamycin resistance gene; RB: left border of T-DNA; RB: right border of T-DNA; T CaMV: terminator from Cauliflower mosaic virus; Pnos: nopalin synthase promoter; nptII, Neomycin phosphotransferase II; (**B**) The severity of symptoms induced in AFCVd-infected (AI) control (CT) and Hl-HM7T lines leaves of hop plants after 10 months of biolistic inoculation of AFCVd. Severe symptoms such as chlorotic mottling, leaf deformation, and stunting were observed in the Hl-HM7T lines compared with the CT - infected hop plants. AFCVd-infected Hl-HM7T lines The plasmid without dimeric AFCVd cDNA construct was used for mock inoculation (MI); (**D**) Dot-blot hybridization analysis of a [^32^P]-dCTP-labeled AFCVd cDNA probe with total nucleic acids isolated from mock inoculated (MI), AFCVd-infected CT, and Hl-HM7T lines leaves of hop; (**D**) Agarose gel electrophoresis analysis of RT-PCR reaction for mock inoculated (MI), AFCVd-infected CT and Hl-HM7T lines leaves of hop after dormancy; (**E**) Strand-specific RT-qPCR profiles of reverse transcribed (+) or (-) strands of mock-inoculated (MI), AFCVd-infected CT and Hl-HM7T lines leaves of hop plants after dormancy. AFCVd strands and their relative concentrations were normalized to 7SL RNA. Each column represents the mean ± S.D. of three replicates of each PCR. Asterisks indicate statistically significant differences, *significant at p<0.05); **significant at p<0.01, *** significant at p<0.001.

### Heterologous expression of HlMyb7 influences the expression of flavonoid biosynthetic genes and growth and development in Tobacco

To further validate the regulatory functions, three independent transgenic T_3_ tobacco lines with higher *HlMYB7* overexpression (Nt-HM7T-2 and Nt-HM7T-5) and lower HlMYB7 overexpression (Nt-HM7T-8) were selected for subsequent analysis (Figure 6a). In contrast to the dark pink flower color of the WT tobacco, these transgenic plants showed stunted growth and a dramatic loss of petal pigmentation, ranging from pale-pink (Nt-HM7T-8) to entirely white color in the Nt-HM7T-2 and Nt-HM7T-5 lines (Figure 6b). Remarkably, the expression of the *HlMYB7* transgene in tobacco correlated with the severity of pigment loss and anthocyanin content in petals (Figure 6c). The expression levels of anthocyanin biosynthesis genes (Figure 6d), chalcone synthase (*CHS*), chalcone isomerase (*CHI*), flavanone-3-hydroxylase (*F3H*), and dihydroflavonol reductase (*DFR*), were significantly decreased in the Nt-HM7T lines compared with WT tobacco (Figure 6e). The reduced expression of these anthocyanin biosynthesis genes correlates with the relative anthocyanin content in Nt-HM7T tobacco lines, implying that *HlMYB7* plays a repressive role in flavonoid biosynthesis and growth and development.

**Figure 6.**
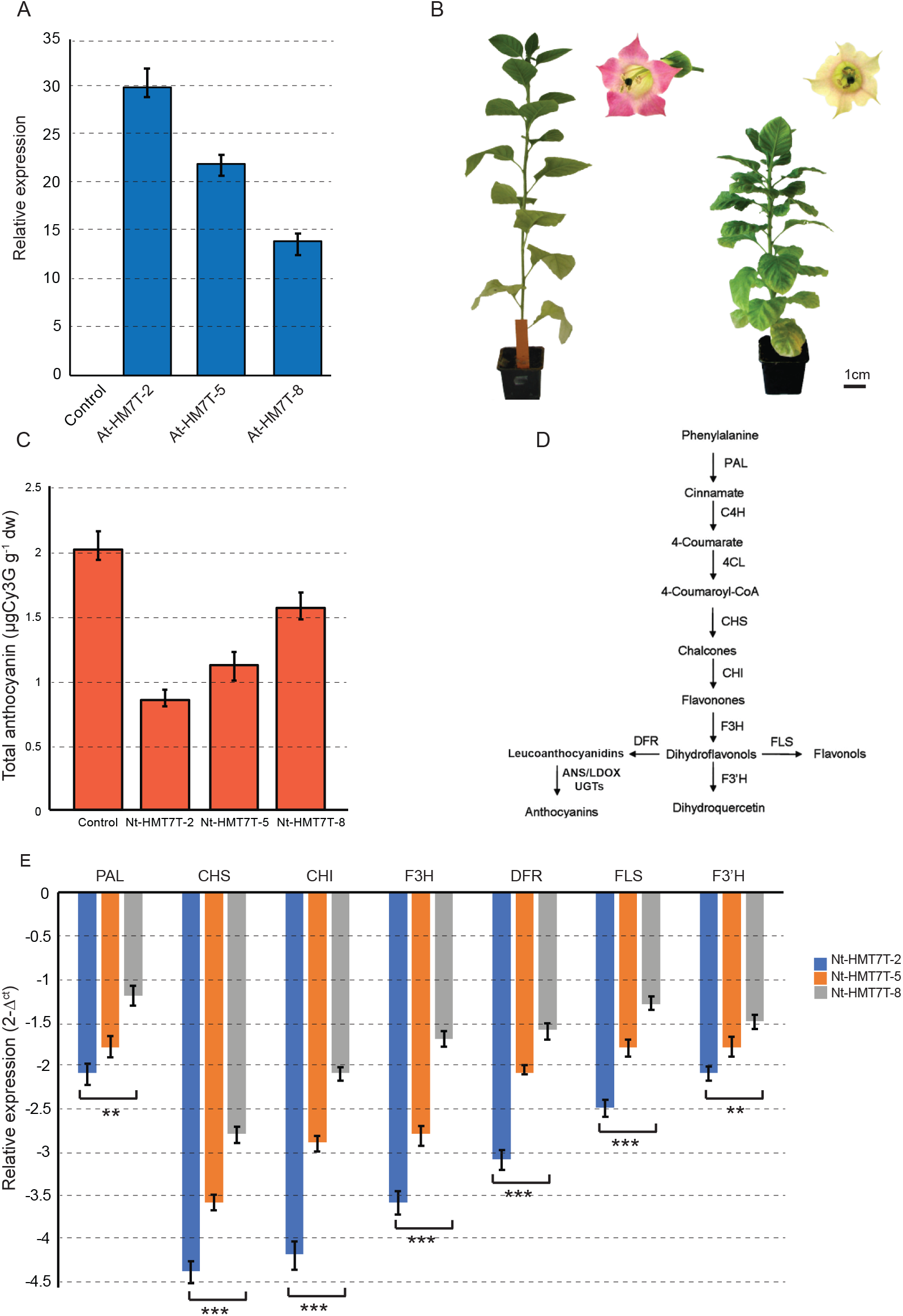
Analysis of the effect of HlMYB7 expression in tobacco plants. (**A**) Relative expression analysis of *HlMYB7* in three representative independent transgenic tobacco lines. The transgenic tobacco carrying the empty vector was used as a control plant. (**B**) Morphological analysis of representative *HlMYB7* overexpression line (Nt-HM7T-2) showing stunted growth and inhibition of flower pigmentation compared to control. (**C**) Total anthocyanin content (µg/g DW) of petal extracts from control and Nt-HM7T tobacco lines. Cyanidin-3-glucoside (*Cy3g*) was used as the anthocyanin standard. Error bars indicate ± SD of three replicates. (**D**) The flavonoid pathway leading to the production of anthocyanins and flavonols in tobacco. The enzymes are as follows: PAL: phenylalanine ammonia lyase, C4H: cinnamate 4-hydroxylase, 4CL: coumarate coenzyme A ligase, CHS: Chalcone synthase, CHI: chalcone isomerase, F3H: flavanone 3-hydroxylase, F’3H: flavonoid 3′-monooxygenase, FLS: flavonol synthase, DFR: dihydroflavonol 4-reductase, ANS: anthocyanidin synthase, LDOX: leucoanthocyanidin di-oxygenase, UGTS: glucose transferase. (E) RT-qPCR analysis of the relative expression levels of flavonoid pathway genes in transgenic tobacco lines overexpressing *HlMYB7* gene. The tobacco *β-actin* gene is used as an internal control, and relative expression levels are determined by the comparative Ct method. Columns represent the average value with SD bar of three technical replicates. Asterisk indicates statistically significant differences, *significant at p<0.05); **significant at p<0.01, *** significant at p<0.001.

### Overexpression of *HlMYB7* influences the expression levels of flavonoid pathway genes and disease severity in *A. thaliana*

To investigate the functional complementarity of *Hl*MYB7 and *At*MYB4, three independent transgenic T3 lines of At-HM7T with higher *HlMYB7* overexpression (At-HM7T-2 and At-HM7T-5) and lower *HlMYB7* overexpression (At-HM7T-6) (Figure 7a) were selected and subjected to qRT-PCR analysis to examine the plausible cross-activation or repression of *A. thaliana* structural genes (*CHS, CHI, F3H, FLS, F’3H* and *DFR*) belonging to the phenylpropanoid pathway (Figure 7b). It is worth noticing that the expression levels of five structural genes of the phenylpropanoid pathway, *CHS, CHI, F3H, FLS, F’3H*, and *DFR* of *A. thaliana* was significantly down-regulated in the At-HM7T-2 and At-HM7T-5 lines compared with At-HM7T-6 (Figure 7c), which correlates with the expression level of the *HlMYB7* transgene in the At-HM7T lines. These findings strongly suggest that *HlMYB7* plays a repressive role on flavonoid structural genes in these plants. Surprisingly, the At-HM7T lines showed reduced axillary branching and stunted growth (Figure 7d), which was in contrast to previous reports that Atmyb4 knockout mutants exhibit the dwarfism phenotype (Wang et al., 2020). Quantitative assessment based on a severity scale of *Plasmodiophora brassiceae* zoospore infection revealed the significant impact of heterologous *HlMYB7* expression on disease severity in *A. thaliana* (Fig. 7e). In *Arabidopsis*, the severity of infection correlates with the expression of HlMYB7, reinforcing its negative role in regulating the defense response and growth and development.

**Figure 7.**
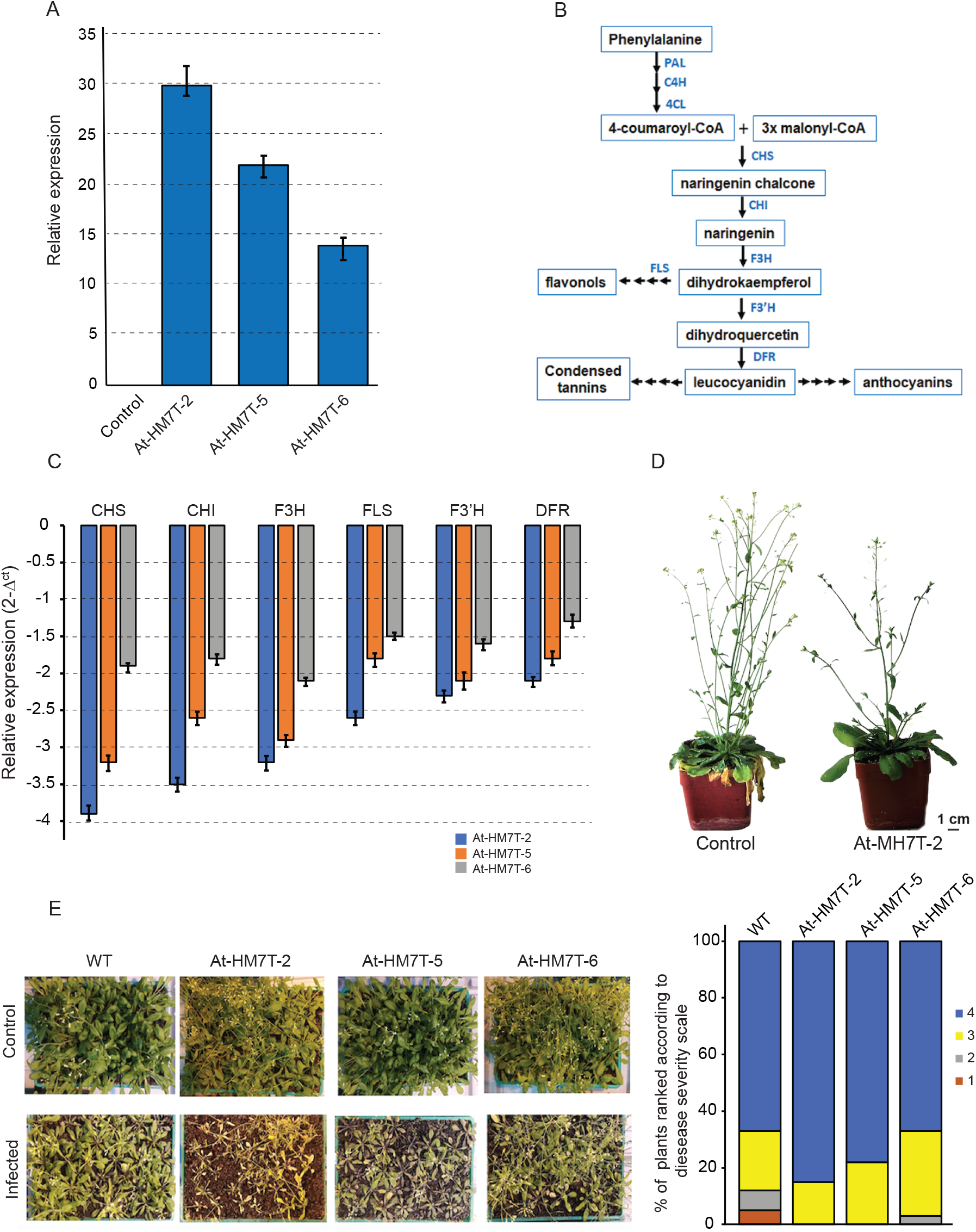
Analysis of the effect of *HlMYB7* expression in *Arabidopsis thaliana* plants. (**A**) Relative expression analysis of *HlMYB7* in three representative independent transgenic *A. thaliana* lines. The transgenic tobacco carrying the empty vector was used as control plant. (**B**) The flavonoid biosynthetic pathway in *A. thaliana*. The enzymes are as follows: PAL: phenylalanine ammonia lyase, C4H: cinnamate 4-hydroxylase, 4CL: coumarate coenzyme A ligase, CHS: Chalcone synthase, CHI: chalcone isomerase, F3H: flavanone 3-hydroxylase, F’3H: flavonoid 3′-monooxygenase, FLS: flavonol synthase, DFR: dihydroflavonol 4-reductase, (**C**) RT-qPCR analysis of the relative expression levels of flavonoid pathway genes in transgenic *A. thaliana* lines overexpressing *HlMYB7* gene. The β-actin gene of *Arabidopsis* is used as an internal control, and relative expression levels are determined by the comparative Ct method. Columns represent the average value with SD bar of three technical replicates. Asterisk indicates statistically significant differences, *significant at p<0.05); **significant at p<0.01, *** significant at p<0.001. (D) Morphological analysis of a representative *HlMYB7* overexpression line (At-HM7T-2) showing stunted growth and rduced axillary branching compared to control. (E) Visual assessment of clubroot disease in WT and At-HM7T *A. thaliana* plants 28 days after inoculation with *P. brassicae*. Disease severity was calculated on a 4-point numerical scale. 0 = no symptoms; 1 = weakly visible symptoms (up to 10% damage); 2 = mild symptoms (10–25% damage); 3 = moderate symptoms (25–50% damage); and 4 = severe symptoms with heavy root galls (>50% damage).

## Discussion

Despite mounting experimental evidence for the involvement of R2R3 MYB proteins as positive and negative transcriptional regulators of structural genes in flavonoid and anthocyanin biosynthesis (Yoshida et al., 2015; Zhai et al., 2016; Rajput et al., 2022), their role in fine-tuning regulatory loops that modulate and maintain PF and BA homeostasis in hop remains ambiguous. In this study, we conducted a systematic study to investigate the regulatory network encompassing the core MBW and WW complexes in the context of feedback regulation, and the role of the R2R3 protein, designated HlMYB7, in the competitive inhibition of these two complexes and structural gene transcripts to design an integral model of the regulation of PF and BA in hop. To this end, we first cloned the *HlMYB7* gene, a homolog of Arabidopsis *MYB4*, and close inspection of the protein sequences revealed that it belongs to MYB subgroup 4 (C4) of the repressor type, which is characterized by the presence of a C1 and C2 domain (EAR repressor) and highly conserved R2 and R3 domains at the N termini. Several transcription factor families contain the C2 repressor motif clade, which can convert a transcriptional activator to a repressor and repress the flavonoid or lignin pathway (Matsui and Ohme-Takagi, 2010), whereas the C4 motif is primarily involved in the negative regulation of volatile benzoid/phenylpropanoid compounds in flowers (Colquhoun et al., 2011). The C1 motif is found in many MYB subgroup 4 proteins, the majority of which are negative regulators of the synthesis of phenylpropanoid-derived compounds (Yoshida et al., 2015). These motifs recruit the chromatin-remodelling complexes and and epigenetically regulate the expression level of enzyme genes (Kagale et al., 2011). The presence of multiple conserved motifs is considered to confer repressive activity to R2R3-MYB repressors of subgroup 4 (Zhou et al., 2019). Several models have been previously implicated to study the interaction of MYB DNA-binding domain (DBD) with the double-stranded DNA, which suggested the cooperative interaction of R2 and R3 with specific DNA consensus sequence (Hichri et al., 2011). In this study, the conserved amino acid signature ([DE]Lx2[RK]x3Lx6Lx3R) of the R3 domain responsible for the formation of a characteristic surface-exposed pattern of hydrophobic and charged residues for interaction with bHLH TF (Zimmermann et al., 2004) was found in the HlMYB7 protein (Figure 1a), suggesting a potential interaction and competition with bHLH TF in binding to structural gene promoters. The amino acid sequence of HLMYB7 shows high sequence identities in its R2R3 domain regions with other R2R3 MYB TFs (Figure 1a), whereas phylogenetic analyses cluster HlMYB7 with other subgroup 4 (SG4) MYB regulators (Figure 1b), which are known to play important roles in phenylpropanoid biosynthesis.

Spatial transcript transcript accumulation analysis revealed the predominant expression of HlMYB7 in lupulin glands (Figure 2a), suggesting that it may play a role in transcriptional regulation of secondary metabolite biosynthesis. Although *HlMYB7* expression was found to be specific to lupulin glands, much lower expression was also observed in other tissues, especially in young leaves, which may be correlated with the presence of lupulin glands on the leaf surface that can biosynthesize secondary metabolites at detectable levels (Mishra et al., 2018). Spatially and temporally restricted expression of MYB proteins involved in the biosynthesis of secondary metabolites has also been reported in other plant species (Cavallini et al., 2015; Rajput et al., 2022). Based on previous findings that MYB-TFs of subgroup 4 can bind to *cis*-acting elements of promoters of structural genes of flavonoid biosynthesis and directly repress their gene expression (Yin et al., 2021), we conducted transient expression studies in leaves of *N. benthamiana*, which showed that HlMYB7 has repressive activity at multiple levels within flavonoid regulatory networks, particularly extreme repression of the key structural gene *HlCHS*_H1 and *HlOMT1*. Intriguingly, it not only directly inhibits the expression of the genes for the biosynthesis of PF and BA, but also the genes encoding the key TFs HlMYB2 and HlWDRI of the MBW and WW activation complexes, respectively (Figure 2). Furthermore, the present study suggests that *HlMYB7* encodes a protein localised in the nucleus that acts as an inhibitor by sequestering the activities of the bHLH and WDR1 proteins of the MBW ternary complex and the WDR1 protein of the WW binary complex (Figure 3). Remarkably, protein-protein interactions revealed that the interaction of HlMYB7 is dependent on bHLH cofactors, highlighting the relevance of the presence of a bHLH interaction motif in the R3 domain of the HlMYB7 protein, which is essential for repressor activity. A number of reports in different plant species have demonstrated the presence of the conserved bHLH interaction motif in R2R3-MYB TFs and their bHLH-dependent repression of flavonol biosynthesis (Xi et al., 2019; Rajput et al., 2022). Thus, the HlMYB7 protein employs a dual lockdown mechanism to reduce flavonoid/flavonol production: first, it prevents the formation of MBW/WW complexes, and then it integrates or binds to these activation complexes and converts them into repressive complexes.

To gain insight into the biological role of *HlMYB7*, we generated constitutive overexpression lines of *HlMYB7* by *Agrobacterium-*mediated transformation. Compared with control plants, the Hl-HM7T lines showed growth retardation, slightly delayed flowering, and susceptibility to viroid infection, but no other significant phenotypic effects. A similar diverse role of R2R3-MYB TFs as negative regulators of phenylpropanoid and flavonoid biosynthesis, growth, and disease resistance has been reported in *Leucaena leucocephala* and *Fragaria ananassa* (Omer et al., 2013; Higuera et al., 2019). Gene expression analysis revealed the impact of overexpression on the remarkable downregulation of PF and BA biosynthetic enzyme gene, including considerable suppression of enzymatic genes of flavonol and anthocyanin metabolic pathways (Figure 4a). The suppresion of PF and BA structural gene expression accompanied by reduction of levels of xanthohumol, alpha and beta acids was observed in transgenic cones and the accumulation pattern of these metabolites coincides with the expression pattern of *HlMYB7* in Hl-HM7T lines (Figure 4b). Nonetheless, we investigated genome-wide changes in steady-state transcript levels using transcriptome profiling to gain a comprehensive understanding of the overall effects of constitutive expression of HlMYB7 in hop. DGE analysis revealed altered gene expression in the Hl-HM7T lines, and it is worth noting that the downregulated genes are involved not only in the PF and BA biosynthetic pathways but also in genes of shikimate metabolism, carbohydrate metabolism, and lipid metabolism (Supplemental Table S2), which may facilitate the diminution of aromatic amino acid supply, precursor and substrate molecules for flavonoid biosynthesis. Substrate channeling is a common event in cellular metabolism, and this type of metabolic reprogramming occurs at the interface of interlocked metabolic enzymes and TF-driven feedback loops (Ma et al., 2018).

Down-regulation of *PAL* transcripts and genes involved in the shikimate pathway and aromatic amino acid biosynthesis in HM7T lines reflects reduced substrate flux towards the flavonoid pathway and supports this trend of regulation and feedback. In addition to the core genes of the flavonoid and BA pathways, genes involved in biosynthesis of terpenoids (terpene synthase 35, beta-amyrin synthase, isopiperitenol/carveol dehydrogenase) and alkaloids (stemmadenine O-acetyltransferase, cytochrome P450 monooxygenases), that play a peripheral role in flavonoid biosynthesis, were also downregulated (Supplemental Table S2). Reduced flavonoid accumulation and decreased expression of disease resistance genes (pathogenesis-related protein: PR-4; PR-5, patatin-like protein 1, thaumatin-like proteins, TIFY proteins, Leucine-rich repeat receptor kinases and ABC transporters, among others) may be attributed to susceptibility of HM7T lines to viroid disease. The involvement of CONSTANS-LIKE (COL), CCT domain-containing genes in early flowering as well as the transcription factor MYC2 and gibberellin-regulated proteins in hormonally controlled developmental steps is well documented in plants (Zhang et al., 2015; Song et al., 2022). Down-regulation of these genes, including a number of genes involved in phytohormone biosynthesis and signaling pathways, may account for the stunted growth and delayed flowering in HM7T lines.

The regulation of anthocyanin and other flavonoid biosynthesis at the transcriptional level is well established in the model plants *A. thaliana*, petunia and tobacco, and therefore serves as a heterologous expression system for understanding the functional conservation of genes as well as their regulatory hierarchy (Kramer, 2015). Heterologous expression of *HlMYB7* reduced stem height and significantly decreased petal pigmentation in Nt-HM7T lines, which was closely correlated with the level of transgene expression and the corresponding decrease in anthocyanin content (Figure 6). We found that key structural genes of anthocyanin metabolism, such as *NtCHI, NtF3’H, NtF3H, NtDFR*, and *NtANS*, were significantly downregulated in the Nt-HM7T lines and showed a negative correlation with *HLMYB7* expression (Figure 6). In this context, it is worth noting that heterologous expression of *HlMYB7* in *Arabidopsis* produced comparable results in terms of down-regulation of key structural genes of flavonoid metabolism, including increased disease susceptibility (Figure 7), supporting its function as a flavonoid repressor and negative regulator of defense response and growth and development. This highlights the complexity and multiple levels of control of HlMYB7 TF in hop. Our findings are consistent with previous studies describing that overexpression of *EgMYB1* resulted in a decrease in lignin/phenylpropanoid content, increased disease susceptibility, and growth retardation in *Arabidopsis* and poplar (Legay et al., 2010).

## Conclusions

In light of our previous and current findings, we propose an ‘accelerator-and-deaccelerator’ model that illustrates the interplay between MBW (HlMYB2/HlbHLH2/HlWDR1) and WW (HlWRKY1/HlWDR1) positive regulator complexes and HlMYB7 as a repressor in fine-tuning the PF and BA pathways (Fig 8). The present study not only advances our understanding of the temporal and spatial regulation of the biosynthesis of PF and BA in lupulin glands, but also opens new avenues for secondary metabolite engineering by using *HLMY7* as a potential candidate gene.

**Figure 8.**
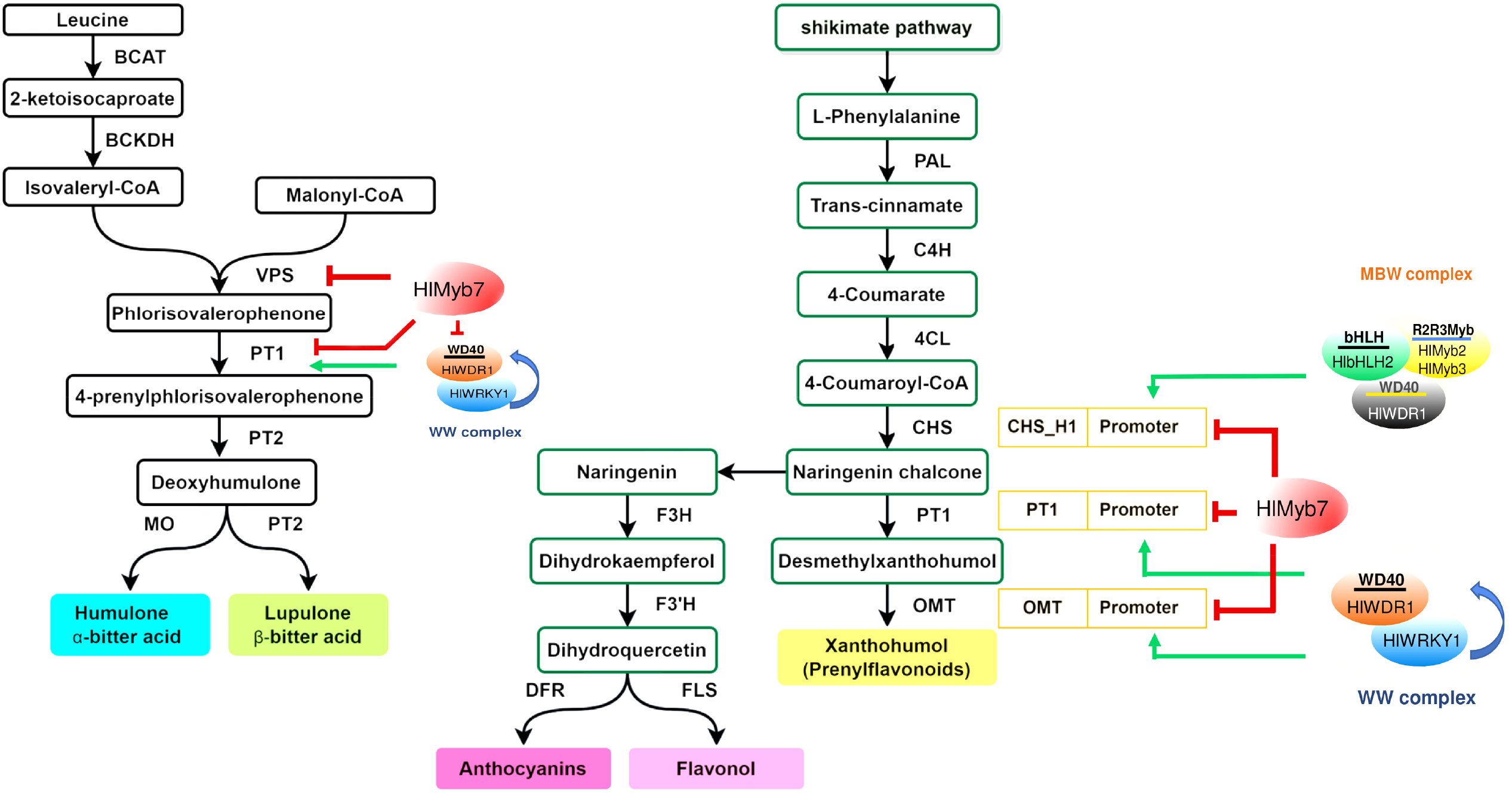
A proposed ‘accelerator-and-deaccelerator’ model for prenylflavonoid (PF) and bitter acids (BA) biosynthesis in hop. The tripartite MBW (HlMYB2/HlbHLH1/HlWDR1) binds to *CHS*_H1 promoter, whereas binary WW protein complexes (HlWRKY1/HlWDR1) bind to the gene promoter of prenyltransferase isoform 1 (*PT1*) and O-methyltransferases (*OMT*) and activate transcription of these genes (marked by the solid green line). In this regulatory network, HlMYB7 can directly repress the transcriptional activity of *CHS*_H1, *OMT, PT1*, and valerophenone synthase (*VPS*) genes (shown by the solid red line) or act as a passive competitor, either interfering with the formation of MBW and WW complexes by regulating their expression or affecting their binding to the promoter sites of the PF and BA genes. HlMYB8 plays a role in competition between flavonol and prenylflavonoid pathways by diverting substrate flux of CHS_H1 gene product to flavonol and anthocyanin pathways

## Material and Methods

### Plant Materials

In this study, hop (*Humulus lupulus* L.) Osvald’s clone 72 was used for *Agrobacterium*-mediated plant transformation and subsequent gene expression experiments. Two-week-old *Nicotiana benthamiana* plants were used for Agro-infiltration-mediated transient expression assays. Naturally occurring *A. thaliana* (ecotype Columbia) and *N. tabacum* (cv. Samsun) were used for transformation and gene expression studies. Plants were grown in a greenhouse environmental control system at a temperature of 25 ± 3 °C under natural lighting conditions with supplemental illumination [170 µmol m^-2^ s^-1^ PAR] to maintain a 16 h diurnal period. Culture conditions for growth of *A. thaliana* and *N. tabacum* seedlings were a 16/8-h light/dark cycle at 23–25 °C with a light intensity of 100 μ mol m^− 2^ s^− 1^ and relative humidity of approximately 50%. In vitro regenerated hop plants were maintained in the growth chamber (Weiss Gallenkamp, UK) at 22 ± 1 °C under 16/8-h light/dark cycle with a light intensity of 200 µmol m^-2^ s^-1^. All collected samples were frozen in liquid nitrogen and stored at −80 °C for further analysis.

### Vector construction, genetic transformation and screening of transgenic lines

The cDNA sequence of *HlMYB7* (GenBank accession number: FR873650) was amplified from our previously constructed lupulin-specific cDNA library of hop cv Osvald’s clone 72 with gene-specific primers containing *Xba*I and *Xho*I restriction sites (Supplemental Table S1). The PCR product was cloned into the pGEM-T vector (Promega, Madison, WI, USA) and confirmed by Sanger sequencing. The insert was cloned into the *Xba*I-*Xho*I sites of the binary vector pLV07 under the control of the cauliflower mosaic virus (CaMV) 35S promoter after digestion with restriction enzymes according to the protocol described previously (Matoušek et al., 2012). The overexpression cassette (35S::*HlMYB7*) was sequenced to confirm the cloning and integrity of the coding sequence *HlMYB7* and subsequently delivered into *A. tumefaciens* strain GV3101 by electroporation method according to the previously described protocol (Holsters et al., 1978). Genetic transformation of hop (cv. Osvald 72) and selection of putative transformants were performed as previously described (Matoušek et al., 2016). The plantlets with 10-15 cm in height with well-developed root systems were transferred to non-sterile conditions (soil with peat moss) and maintained in the growth chamber (16 h daylight, 5000 lx, 22–24 °C) for 4 to 5 weeks for acclimatization prior to transfer to a greenhouse and further to an outdoor containment facility meeting the necessary requirements for handling GMOs. Molecular confirmation of T-DNA insertion events in transgenic lines was performed by genomic PCR, real time RT-PCR analysis using gene-specific primers (Supplemental Table S1), and Southern blot hybridization, as described previously (Tai and Tanksley, 1990). Visual observations were made at regular intervals to assess the growth and morphological characteristics (height, number of nodes, density of lupulin glands, and leaf morphology) of the Hl-HM7T lines (1, 2, and 11) compared with the control. In addition, the chlorophyll content of the Hl-HM7T lines and control hop regularly evaluated by recording the absorbance of their leaf extract at 646 and 663 nm wavelengths using UV spectrophotometry (Shimadzu, Japan) according to the method described previously (Lichtenthaler and Wellburn, 1983).

Transfomation of *A. thaliana* was performed using the floral dip method (Clough and Bent, 1998) with the constructs of interest in *A. tumefaciens* (35S::*HlMYB7*). The homozygous T3 single insertion lines of At-HM7T were screened on media containing kanamycin (50 μg/ mL). Leaves of sterile shoot cultures of *Nicotiana tabacum* (cv. samsun) were used for transformation with *A. tumefaciens* harboring 35S::*HlMYB7* construct according to the leaf disk transformation and regeneration protocol as previously described (Rosahl et al., 1987). The kanamycin-resistant T_0_ generation of tobacco lines was transferred to a growth chamber for rooting and acclimation to harvest seeds of the T_1_ generation. The transgenic T_3_ homozygous Nt-HM7T lines were used for subsequent experiments.

### Phylogenetic and sequence analyses

The full-length amino acid sequences of the known R3 MYB repressors were retrieved from the NCBI (National Centre for Biotechnology Information) public databases. For the construction of the phylogenetic tree, multiple alignments of the full-length protein sequences were performed with the ClustalW algorithm of the MEGA software package (version 10.1.8; Pennsylvania State University, PA, USA) (Kumar et al., 2008) using the default parameters. All positions with alignment gaps and missing data were eliminated before generating the phylogenetic trees. A maximum likelihood phylogenetic tree was constructed and tree nodes were scored using the bootstrap method for 1000 replicates. Analysis of conserved motifs in the multiple sequence alignment of R2R3-MYB TFs was performed using MEME Suite software (http://meme-suite.org/) (Bailey et al., 2009), with parameters set to a maximum of five motifs and zero or one occurrence of the distribution of motif sites per sequence.

### Transcriptional activation assay in *N. benthamiana* leaves

Transcriptional activation of HlMYB7 at the promoters of the PF and BA pathway genes was analysed by the fluorometric β-glucuronidase assay (GUS) in transiently transformed leaves of *N. Benthamiana*, as described previously (Chen et al., 2005). The full-length coding sequence (CDS) of *HlMYB2, HlMYB7, HlWDR1, HlbHLH2, HlWRKY1, HlMYB8* was amplified with gene-specific primers (Supplemental Table S1) and cloned into the plant binary vector pLV-07 under the CaMV35S promoter. The promoter fragments 1.5 kb upstream regions from the start codon of the structural genes (*HlCHS*_H1, *HlOMT1, HlVPS*, and *HlPT*-1) and TF-encoding genes (*HlMYB2, HlMYB7, HlWDR1, HlbHLH2, HlWRKY1*, and *HlMYB8*) of the PF and BA biosynthetic pathway of hop were cloned into the pBGF-0_GUS vector using the primers (Supplemental Table S1). These constructs were individually transformed into *Agrobacterium tumefaciens* strain LBA4404 using the electroporation method, and transient expression analysis of β-glucuronidase (GUS) was performed as described previously (Matoušek et al., 2016).

### Subcellular Localization of *HlMYB7*

The full-length CDS of *HlMYB7* was amplified with gene-specific primers (Supplemental Table S1) from cDNA libraries and ligated into the *Xho*I sites of binary vector pEarleyGate 103 to construct the CaMV 35s: *HlMYB7-GFP* plasmid. After confirmation of sequencing, CaMV 35s: *HlMYB7-GFP* plasmid and empty vector were infiltrated into the epidermal cells of *N. benthamiana* leaves as described previously (Liu et al., 2010). The GFP fluorescence signal was visualized and photographed using a confocal laser scanning microscope (Leica TCS SP5, Germany).

### Yeast two-hybrid assays

For yeast two-hybrid (Y2H) screening, the full-length coding sequence of *HlbHLH2, HlMYB2, HlMYB8, HlWDR1*, and *HlWRKY1* was individually cloned into the bait vector pGBKT7-GW [DNA-binding domain (BD)] (Addgene Inc, MA, USA) to produce fusion proteins containing a GAL4 activation domain (AD; prey), while the full-length coding sequence of *HlMYB7* was cloned into the prey vector pGADT7-GW [DNA-binding domain (BD)] (Addgene Inc, MA, USA) using the In-Fusion Snap Assembly Kit (Takara Bio). The bait and different combinations of prey vectors were cotransformed into yeast strain AH109 using a modified yeast transformation protocol (James et al., 1996). The transformed yeast cells were plated out on selective medium lacking Trp, Leu, and with Ade and His (SD/-Trp/-Leu/+Ade/+His) and allowed to grow for 3 days at 30 °C. Subsequently, colonies were inncoulated into Broth medium (SD/-Trp/-Leu/+Ade/+His) and allowed to grow overnight at 30 °C. Growth of colonies was maintained at 0.3 optical density (OD) for each spotted 5 µl with decreasing concentraions of 1, 10^−1^, 10^−2^, 10^−3^ on minus Ade, His (SD/-Trp/-Leu/-Ade/-His) and with sprayed 100 µl of 4 mg/ml X-α-Gal (Takara Bio, CA, USA) in dimethylformamide on plates (SD/-Trp/-Leu/-His/X-α-Gal). Co-transformation of empty prey and bait construct and empty bait with prey construct were used as controls.

### Differential gene expression analysis and Functional categorization

DNase-treated total RNA isolated from leaves of *HlMYB7-*overexpressing transgenic lines (Hl-HM7T-1, Hl-HM7T-2, and Hl-M7T-11) and CT hop plants, was quantified using the NanoDrop 2000 spectrophotometer (Thermo Scientific, Waltham, MA, USA), and integrity was determined using the RNA 6000 Nano Assay Kit (Agilent, USA) in the Agilent 2100 Bioanalyzer (Agilent, California, CA, USA). Isolation of mRNA was performed with the polyATract mRNA Isolation System IV (Promega Corporation, Madison, WI, USA) and was further processed after quality assessment and enrichment for pair-end cDNA library construction using the TruSeq RNA Library Preparation v2 Kit (Illumina, USA) according to the manufacturer’s instructions. The quality of the libraries was assessed using the Agilent 2100 High Sensitivity DNA Kit (Agilent Technologies, Santa Clara, CA, USA) and subjected to paired-end sequencing on the Illumina HiSeq 2500 platform (Illumina, San Diego, CA, USA). Raw sequencing data were processed using fastp software in order to remove adapters, low-quality reads, k-mers contamination, and low-quality regions (Chen et al., 2018). The quality of reads was evaluated based on their error rate, GC content bias, and base quality scores (phred score quality below 30) using the FASTQC toolkit (Andrews, 2010), and 50 bp or longer paired-end reads were selected for downstream analysis. The obtained high-quality, clean reads of each sample were mapped to the full length transcripts of hop (HopBase genomic resources repository; http://hopbase.org/) using StringTie software (Pertea et al., 2015). The ambiguously aligned reads were discarded, and the uniquely located reads were used to calculate read counts for each gene. The FPKM value (fragments per kilobase of transcript per million mapped reads) was determined for each sample using the expectation maximization method (RSEM) and used for alignment-based abundance estimation (Li and Dewey, 2011). Read counts for each transcript were analyzed and used for differential expression analysis, the logarithmic fold change, and false detection rate (FDR) for each transcript using the Bioconductor software package DESeq2 (Love et al., 2014) with default parameters. In this study, unigenes with an FDR of less than 0.05 and an absolute value of a log_2_-fold change greater than 2-fold change between samples were considered differentially expressed genes (DEGs).

### Functional annotation and biological classification of differentially expressed genes

Identified differentially expressed genes (DEGs) were subjected to BLASTX analysis against NCBI nonredundant (nr) proteins database using an *E*-value cut-off of 1.0 E^−5^, and the Blast2GO tool v.4.1 (www.blast2go.com) (Conesa et al., 2005) was used to obtain GO annotations and functional classifications. GO terms enriched in DEGs were identified using Gene Set Enrichment Analysis (GSEA) of the clusterProfiler Bioconductor package (Yu et al., 2012). The web-based annotation server KAAS (KEGG Automatic Annotation Server; version 2.1) (www.genome.jp/kegg/kaas) was used to assign KEGG (The Kyoto Encyclopedia of Genes and Genomes) pathways to DEGs using the single-directional best-hit (SBH) method with a default threshold of 60-bit score (Moriya et al., 2007). Gene set enrichment analysis (GSEA) was performed using the online tool WebGestalt to investigate the statistical enrichment of DEGs in KEGG pathways (Liao et al., 2019). The functional terms and pathways were considered significantly enriched at Bonferroni-corrected P-values ≤ 0.05. Furthermore, DEEGs were mapped to different functional categories with the Mercator tool (https://www.plabipd.de/portal/mercator-sequence-annotation) using default parameters, and mapping file was imported into MapMan software (version 3.6) for visualization of gene expression data (Thimm et al., 2018).

### RNA extraction and quantitative real-time PCR analyses

Total RNA from different tissues of hop, *N. tabacum, A. thaliana* control, and transformed lines was extracted using Concert™ Plant RNA Purification Reagent (Invitrogen, USA). Removal of genomic DNA contamination and first-strand cDNA synthesis were performed using the Ambion® DNA-free™ DNase Kit (Thermo Fisher Scientific, USA) and the Superscript® III First-strand cDNA Synthesis System (Invitrogen, USA), respectively, according to the manufacturer’s instructions. Quantitative real-time PCR (qRT-PCR) experiments were performed using diluted cDNA (1:50) with 200 nM of each gene-specific forward and reverse primer (Supplemental Table S1) mixed with SYBR^®^ Green supermixes (Bio-Rad, USA) according to the manufacturer’s instructions on a Thermal Cycler CFX96 Real-Time System (Bio-Rad, USA) to detect the changes in the expression patterns of genes associated with the biosynthesis of PF and BA in hop and flavonoid biosynthesis in *A. thaliana* and *N. tabacum*. Data were analyzed and quantified using Bio-Rad IQ™5 Optical System version 2.0 software, and the product specificity size was confirmed by melting analysis. Similarly, the reliability of RNA-seq results was confirmed by qRT-PCR analyzes of ten randomly selected DEGs (five up-regulated and five down-regulated genes) and genes related to flowering time, immune response, and growth and development. The expression of glyceraldehyde-3-phosphate dehydrogenase in hop (ES437736) and actin gene of *N. tabacum* (JQ256516.1), *A. thaliana* (AT3G18780) was used to normalize the different Ct values. The relative expression level of the genes was calculated using the comparative Ct method (2^−ΔΔCt^) as described previously (Livak and Schmittgen, 2001). All analyses were performed in triplicate and repeated twice with three biological replicates. All statistical analyses were performed using the SAS software package and the Duncan multiple range test (Duncan, 1955).

### Assessment of plant disease resistance

To investigate the effects of *HlMYB7* overexpression on disease severity, leaves of six-month-old Hl-HM7T lines and control hop were artificially infected five times with 250 ng of a monomeric cDNA-infectious construct of *Apple fruit crinkle viroid* (AFCVd), as previously described (Matoušek et al., 2017). Immediately after inoculation, plants were covered with polyethylene bags to prevent desiccation of the shot-wound leaf area and placed in darkness for 24 h. Subsequently, the mock- and AFCVd-inoculated plants were cultivated in their natural environment and visually examined for symptom development. Systemically infected and mock-inoculated leaves were harvested 270 days post inoculation (dpi) after completion of dormancy from the shoot apex after appearance of typical symptoms of AFCVd-infection (Matoušek et al., 2017). Strand-specific RT -qPCR (ssRT-qPCR) and dot-blot analysis were performed to monitor the titer of AFCVd infection as previously described (Matoušek et al., 2017).

Fourteen-day-old *A. thaliana* control and At-HM7T seedlings grown in a controlled environment (23°C, 16-h light, 100 mmol photons/s/m2) using a compost-sand mixture (9:1 v/v) (pH 5.8) were inoculated with 1 ml dormant spore suspension of *Plasmodiophora brassiceae* adjusted to 10^4^ to 10^5^ colony forming units (cfu) ml^-1^ in Na_2_HPO_4_ buffer (pH 5.8) into the soil around each plant. Control plants were grown under the same conditions and treated with Na_2_HPO_4_ buffer (pH 5.8) instead of the spore suspension (mock). The severity of infection was visually scored 28 days after inoculation on a 5-point numerical scale based on the percentage infected plants (Ludwig-Müller et al., 2015): 0 = no symptoms; 1 = weakly visible symptoms (up to 10% damage), 2 = mild symptoms (10–25% damage); 3 = moderate symptoms (25–50% damage); and 4 = severe symptoms showing strong root galls (>50% damage). Disease index (DI) was calculated according to Leath et al. (1989) using the following equation DI = (1n_1_ + 2n_2_ + 3n_3_ + 4n_4_)100/4N_t_, where n_1_-n4 is the number of plants in the indicated classes and N_t_ is the total number of plants examined. For each biological experiment, 30 *A. thalaina* plants were analysed. Means and SE were determined with Fisher’s LSD test (α = 0.05) using Statistix software (v9.0).

### Metabolite analysis

Dried cones were harvested from control and Hl-HM7T lines and freeze-dried before determination and quantification of hop resins (α- and β-acids) and polyphenols (PFs). Identification and quantification of hop resins, BAs, xanthohumol (XN), and demethylxanthohumol (DMX) were performed using the HP Agilent 1100 high-performance liquid chromatography (HPLC) system (Hewlett-Packard, Waldbronn, Germany) from lypholized tissue as previously described (Mishra et al., 2018). HPLC analyses and quantification of total anthocyanins in transgenic and control tobacco flowers were performed according to the protocol described previously (Deluc et al., 2006).

### Accession numbers

Sequencing data used in this study were deposited to NCBI SRA under the submission numbers: SRX17180127 (CT1), SRX17180128 (CT2), SRX17180129 (CT3), SRX17180130 (Hl-HM7T-1), SRX17180131 (Hl-HM7T-5) and SRX17180132 (Hl-HM7T-11).

## Supplemental data

**Supplemental Figure S1**. Molecular analysis of transgenic hop lines overexpressing HlMYB7 (**A**) Schematic representation of the T-DNA region of the plant expression vector pLV07 used for hop transformation. RB and LB: right and left T-DNA borders, Pnos: nopaline synthase promoter, nptII: neomycin phosphotransferase II gene, P35S, cauliflower mosaic virus 35S promoter; TCaMV, cauliflower mosaic virus terminator. (**B**) Southern hybridization analysis of hop plants carrying T-DNA with HlMYB7 transgene. Genomic DNA of individual kanamycin-resistant hop lines was digested with *EcoR*I and hybridized with 0.8 kb 32P[dCTP]-labeled cDNA probe. The membrane was visualized by phosphorimaging using a Typhoon 9200 imager (Amersham Pharmacia). Internal HlMYB7 gene is shown by an arrow. The 32P[dCTP] labeled bands of the 24 kb DNA ladder (Fisher BioReagents™) are shown on the left side of the gel. (**C**) Gel image of the semi-quantitative RT-PCR assay of HlMYB7 in transgenic hop leaves carrying HlMYB7 gene (**D**) Relative expression analysis of HlMYB7 in seven independent transgenic hop lines. The transgenic hop carrying the empty vector was used as a control plant (**E**) Phenotypic comparison of the growth of in vivo-grown six-month-old HLMYB7-overexpressing and control hop plantlets. **Supplemental Figure S2**. (A) GO enrichment analysis of differentially expressed genes in leaf of HlMYB7 overexpressing lines compared with control at a significance level of p = 0.05, using GO terms from GO Slim. The x-axis labels represent fold enrichment, and the y-axis labels represent GO terms. The color gradient represents the adjusted P values, and the differences in bubble size correlate with the enrichment factor. Only the categories showing statistically significant enrichment at the gene expression level. (B) KEGG pathway enrichment analysis for differentially expressed genes in leaves of HlMYB7-overexpressing lines compared to control at a significance level of p = 0.05. (C) MapMan visualization of significantly differentially expressed transcripts (log_2_ fold change) associated with hormone signaling, transcription factors, ROS signaling processes, and secondary metabolites in HlMYB7-overexpressing lines compared to control. Red and green signals in the scale indicate up-regulation and down-regulation of transcripts, respectively, in Hl-HM7T lines compared to control hop plants.

**Supplemental Figure S3**. Validation of RNA sequencing data by RT-qPCR. (A) The results of qPCR validation of top five upregulated [DNA damage-repair/toleration protein (DRT), Histone 4 (HIS4), calcium-binding protein CML37 (CBP), small nuclear ribonucleoprotein (SNR), and photosystem I assembly protein ycf3 (YCF3)], top five downregulated [beta-galactosidase-like isoform X1 (BGAL), TLC domain-containing protein (TDP), Cysteine proteinase (CP), asparagine synthetase (AS) and mulatexin (MUL)], including CONSTANS-LIKE (COL) and CCT domain-containing genes, patatin-like protein 2 (PP2), pathogenesis-related protein 4 (PR-4), and TIFY proteins. (B) The results of qPCR validation of genes constant6 (CON), patatin-like protein 1 (PAT), pathogenesis-related thaumatin-like protein (PRP), and chitinase class IV (CHT) genes. Log2 fold change (Log2FC) change of each gene was calculated by the 2−ΔΔCT method and compared with the results of RNA-seq (grey column). The log_2_ fold change (Log_2_FC) of each gene was calculated using the 2^-ΔΔCT^ method and compared with the RNA-seq results (grey column). The graph shows values ± SD of three leaves of Hl-HM7T-1, Hl-HM7T-5, and Hl-HM7T-11 transgenic lines of hops.

**Supplemental Table S1**. Primers used for cloning into the plant vector, probe preparation, and qRT-PCR analyses.

**Supplemental Table S2**. Morphological and physiological characteristics of the Hl-HM7T lines compared with control hop plant.

**Supplemental Table S3**. Unigenes differentially expressed between leaves of control and transgenic HlMYB7 plants of hops.

## Acknowledgments

We acknowledge the technical supports recieved from Ganesh Selvaraj Duraisamy, Helena Matoušková, Olga Horáková, Lidmila Orctová, Kavita Mishra and Jana Jehlíková (Institute of Plant Molecular Biology, BC-Czech Academy of Sciences Centre). We are grateful to. Petr Svoboda (Hop Research Institute Co Ltd. Žatec, Czech Republic) for technical assistance in the maintenance of transgenic hop plants. The authors declare no competing financial interests.

## Author contributions

A.K.M., T.K., J.M., and K.M.A.A. conceived the research plans and designed the research. A.K.M., T.K., V.S.N., A.K., and J.M. were all involved in the experimental work. KK performed phytochemical analysis. N.S. and K.M.H. performed bioinformatics analyses. A.K.M., and T.K. wrote the manuscript. J.M. and K.M.A.A. revised the manuscript. All authors read and agreed to submit the manuscript for publication.

